# Non-Canonical Activation of HSF1 Stimulates mTORC1-Mediated Translation in HCMV-Infected Monocytes

**DOI:** 10.64898/2026.07.21.739842

**Authors:** Michael J. Miller, Shima Moradpour, Brittany W. Geiler, Jamil Mahmud, Wayne A. Decatur, Gary C. Chan

## Abstract

Human cytomegalovirus (HCMV) is a major cause of organ disease among immunonaïve and immunocompromised individuals. HCMV infection stimulates the survival of normally short-lived circulating monocytes, allowing these blood cells to mediate the dissemination of the virus from the initial point of infection to distant organ sites. We previously showed that HCMV induces a non-canonical phosphorylation of Akt within infected monocytes that activates the stress response transcription factor Heat Shock Factor 1 (HSF1). In this study, we demonstrate that HSF1 is necessary for the survival of HCMV-infected monocytes using both pharmacological and genetic approaches. In contrast, HSF1 inhibition had minimal effect on the viability of uninfected cells, indicating the specific involvement of HSF1 on the survival of infected monocytes. Surprisingly, the aberrant activation of HSF1 by HCMV did not trigger nuclear relocalization, suggesting that HSF1’s regulation of monocyte viability occurs within the cytoplasm. Indeed, we found that HCMV-activated, cytoplasmic HSF1 directly binds to mTOR, a critical component of the mTORC1 complex involved in the regulation of mRNA translation. SUnSET (Surface Sensing of Translation) assays revealed HCMV-activated HSF1 increases mRNA translation through mTORC1. Ribosomal profiling identified the increased translation of a selected subset of pro-survival transcripts, including cIAP2, which we validated to selectively stimulate the survival of HCMV-infected monocytes. Taken together, these data demonstrate that the non-canonical activation of HSF1 in infected monocytes drives mTORC1-dependent translation of antiapoptotic transcripts, ensuring the survival and dissemination of infected monocytes.

**IMPORTANCE:** HCMV is a primary driver of morbidity and mortality in individuals with compromised or immature immune systems. Spread of HCMV throughout the body relies on the infection of peripheral blood monocytes, which spread the virus to end-organ tissues. However, the naturally short lifespan of monocytes must be overcome to allow for viral spread to occur. Here, we demonstrate that HCMV uniquely regulates the cellular stress response to promote the long-term survival of infected monocytes. Specifically, HCMV activates the cellular stress response transcription factor HSF1 to block the progression of apoptosis. In contrast to traditional heat shock stress where HSF1 translocates into the nucleus to mediate transcription, HCMV infection retains activated HSF1 in the cytoplasm where it binds to mTOR to promote protein synthesis of prosurvival factors necessary for the survival of infected monocytes. Overall, our study provides insight into the complex regulator mechanisms through which HCMV usurps host stress responses to promote viral dissemination.

## INTRODUCTION

Human cytomegalovirus is a ubiquitous betaherpesvirus with an estimated seroprevalence between 50-100% globally [1–3]. HCMV infection in healthy immunocompetent individuals are often self-limiting, although it may present as mild mononucleosis-like symptoms [4–6]. HCMV infection is also associated with increased risk of several inflammatory disorders, including atherosclerosis and restenosis, as well as cancers, including breast, colon, and glioblastoma cancers [7–11]. In contrast, HCMV is a significant cause of morbidity and mortality for patients with compromised and immature immune systems, including those undergoing chemotherapy, immunosuppression in preparation for transplantation, HIV/AIDS patients, and neonates [12–17]. Acute HCMV disease is often characterized by widespread viral dissemination to multiple organ sites and severe inflammatory end-organ damage [6, 18, 19].

Circulating monocytes are thought to mediate the systemic dissemination of HCMV as they are the primary cell type infected in the blood and the predominant infiltrating cell type found in the organs of transplant recipients [20–24]. However, monocytes are inherently short-lived cells in the absence of survival stimuli with a lifespan of ∼48 hours (h) following release from the bone marrow [25, 26]. Furthermore, monocytes are not permissive for HCMV lytic replication, making these blood cells ill-suited to facilitate the spread of infectious progeny [27, 28]. To circumvent these shortcomings, we previously showed that HCMV binding and entry stimulate the survival and differentiation of short-lived monocytes into long-lived macrophages permissive for viral replication [29–31]. Mechanistically, HCMV induces a non-canonical phosphorylation of Akt leading to the activation of mTORC1 [32–36], which stimulates the translation of antiapoptotic transcripts without inducing the translation of several antiviral transcripts within HCMV-infected monocytes [32, 34, 36, 37]. The increased synthesis of mTORC1-dependent pro-survival factors supports the survival and differentiation of HCMV-infected monocytes, thereby facilitating the spread of the virus following a primary infection [36].

The mTORC1 complex serves as a major regulatory signaling hub controlling cap-dependent protein synthesis [38–42]. In contrast to aberrant HCMV-activated Akt, canonically activated Akt induced by normal myeloid growth factors does not induce mTORC1 activity [36], indicating a unique viral mechanism responsible for stimulating mTORC1-dependent translation within infected monocytes. During times of cellular stress, including viral infections, mTORC1 is typically inhibited in an AMPK-dependent fashion. AMPK ensures the integrity of the TSC1/2 complex that regulates cell growth by inhibiting mTORC1 [43–45]. During lytic HCMV infection, the negative relationship between AMPK and mTORC1 is ablated through the action of the viral protein pUL38 [46, 47]. Similarly, we previously showed that HCMV also sustains mTORC1 activity, and subsequent translation of cap-dependent pro-survival mRNAs, in quiescently infected monocytes despite the presence of AMPK [34, 36]. However, the absence of pUL38 expression in quiescently HCMV-infected monocytes indicates a distinct mechanism responsible for uncoupling AMPK and mTORC1 activity. Indeed, we previously demonstrated that the stress responsive cellular transcription factor Heat Shock Factor 1 (HSF1), which directly interacts with and represses AMPK [48], is necessary to maintain mTORC1 activity within infected monocytes [34].

HSF1 is a ubiquitous stress-responsive transcription factor that regulates the gene expression of several of heat shock proteins (HSPs), a family of chaperones responsible for protecting cells against cellular stresses [49, 50]. Under homeostatic conditions, HSF1 exists in an inactive monomeric state bound to an obligatory heat shock protein chaperone, most often HSP70 or HSP90 [49, 51]. Following an insult to the cell, HSF1 and its chaperone quickly dissociate allowing for HSPs to facilitate proper protein folding, repair damaged proteins, or degrade those beyond repair to limit proteotoxic stress within cells [49, 52, 53]. Simultaneously, unbound HSF1 trimerizes and enters the nucleus, binding to heat shock elements (HSEs) upstream of target gene promoters [54, 55]. Once bound to HSEs, HSF1 aids in the recruitment of RNA polymerase II, p-TEFb, and DSIF to initiate transcription of stress-responsive genes [50, 56–58]. Although HSF1’s role in gene regulation is clear, the cellular effects of HSF1 activation during times of cellular stress is more complex than simple regulation at the level of transcription. The HSF1 interactome profoundly transforms in response to stress with the acquisition of new nuclear and cytoplasmic protein partners, indicating that activated HSF1 has diverse cellular functions throughout the cell. Monomeric HSF1 engages several cytoplasmic proteins during stress, including AMPK, Akt, mTOR, MEK and ERK, to regulate a broad array of cellular functions from metabolism to translation [59, 60].

In this study, we report that HCMV rapidly phosphorylates and maintains HSF1 activity in infected monocytes, which contrasts with the transient activation driven by canonical heat stress (HS). This aberrant activation of HSF1 is necessary for the survival of infected monocytes. Consistent with our previous study, HCMV-induced HSF1 promotes mTORC1 activity, driving the expression of several antiapoptotic proteins[36]. We now show that HSF1 binds to mTOR, a component of the mTORC1 complex, suggesting that cytoplasmic HSF1 regulates mRNA translation within quiescently infected monocytes. In support of this, HCMV-activated HSF1 is retained in the cytoplasm of quiescently infected monocytes while HS-activated HSF1 rapidly translocates to the nucleus. Additionally, we found HSF1 inhibition attenuates translation stimulated within HCMV-infected monocytes but had no effect on translation induced by myeloid growth factors or during lytic infection of fibroblasts. Polyribosomal profiling analysis identified the translation of several overlapping antiapoptoic transcripts dependent on both HSF1 and mTORC1, including cIAP2. Inhibition of cIAP2 led to the cell death of HCMV-infected monocytes while having minimal effect on the viability of uninfected cells. Taken together, our results suggest that HCMV utilizes the cytoplasmic HSF1 to sustain translation of pro-survival factors, via its interaction with mTORC1, leading to the long-term survival of short-lived monocytes.

## RESULTS

### HCMV stimulates a distinct HSF1 activation in infected monocytes

We have previously shown that HCMV rapidly induces a sustained phosphorylation of HSF1 during lytic infection of fibroblasts [61]. To determine the kinetics of HSF1 activation during a quiescent infection, primary monocytes were infected with HCMV strain TB40E for 30 minutes (min) and 24 h and HSF1 phosphorylation at Ser326, a hallmark of HSF1 activation, and total HSF1 protein levels measured (Fig. 1A). We found that HCMV infection of monocytes stimulated an early activation of HSF1 at 30 min post-infection (mpi) that was sustained through 24 h post-infection (hpi). In contrast, heat shock (HS) treatment induced a transient HSF1 phosphorylation, which was completely resolved by 24 post treatment (Fig. 1A). As we have previously demonstrated [34], HSF1 activation following HCMV entry is dependent on PI3K/Akt signaling and to a lesser extent, on mTOR (Fig. 1B). Here, we further show that the activation of HSF1 by HS treatment does not require signaling from PI3K/Akt/mTOR. Taken together, these data demonstrate that HCMV infection rapidly stimulates a non-canonical HSF1 activation in infected monocytes that is kinetically and mechanistically distinct from the classical heat shock response.

**Figure 1:**
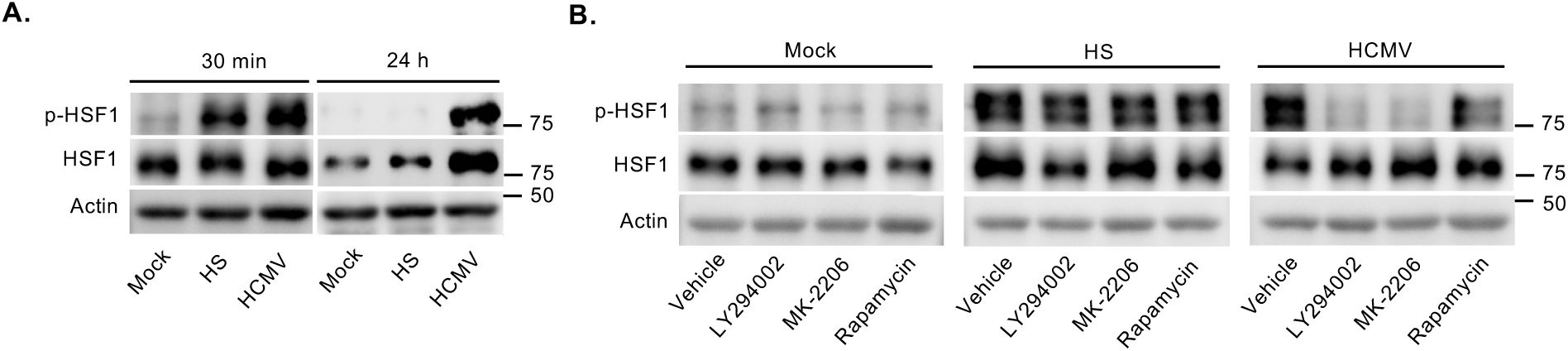
HCMV stimulates a distinct HSF1 activation kinetics following infection of monocytes. **(A)** Primary peripheral blood monocytes were mock- or HCMV-infected (MOI of 5) for 30 min or 24 h. As a positive control, monocytes were subjected to heat shock (HS) for 30 min at 42°C. **(B)** Monocytes were pretreated with LY294002 (a PI3K inhibitor), MK-2206 (an Akt inhibitor), or Rapamycin (a mTOR inhibitor). Cell were then HS treated or HCMV infected for 30 min. Total HSF1 and phosphorylation of HSF1 at Ser326 (p-HSF1) were detected by Western blot. Actin was used as a loading control. Western blots are representative of at least three biological replicates per group.

### HSF1 promotes the survival of HCMV-infected monocytes

To investigate the role of non-canonically activated HSF1 during HCMV infection, we utilized two small molecule inhibitors of HSF1: SISU-102, which directly binds to the amino-terminal HSF1 DNA-binding domain to promote ubiquitin-proteasome-mediated degradation of active nuclear HSF1 [62, 63], and KRIBB11, which binds to the transactivation domain of HSF1 preventing the recruitment of transcription machinery [64]. Monocytes were pretreated with SISU-102 or KRIBB11 for 1 h prior to a 24 h infection with HCMV. As expected, SISU-102 reduced the abundance of HSF1, leading to a significant reduction in phosphorylated HSF1 in HCMV-infected monocytes (Fig. 2A). Consistent with its mechanism of action, KRIBB11 had minimal effect on both the phosphorylated and total levels of HSF1 induced by HCMV. Other reports have demonstrated that inhibition of HSF1 by KRIBB11 reduces the abundance of Mcl-1, an antiapoptotic member of the Bcl-2 family of proteins [65]. Accordingly, suppression of HSF1 by SISU-102 prior to HCMV infection reduced the protein levels of several antiapoptotic proteins, including Mcl-1, HSP27, and XIAP, that we have previously shown to be induced and necessary for the survival of HCMV-infected monocytes [34, 66] (Fig. 2B). In agreement and further validating the on-target effects of SISU-102, knockdown of HSF1 with siRNA, which resulted in ∼90% knockdown, also significantly reduced Mcl-1, HSP27, and XIAP abundance in HCMV-infected monocytes (Fig. 2C). Next, we sought to uncover the role of HSF1 on the viability of HCMV-infected monocytes. The lethal dose 50 (LD50) of uninfected versus HCMV-infected monocytes was determined by treating cells with increasing concentrations of SISU-102, followed by flow cytometric analysis of annexin V and propidium iodide (PI) positive cells. We found that HCMV-infected monocytes are more sensitive to SISU-102-induced cell death, with the LD50 shifting from ∼20 μM in uninfected monocytes to 5.8 μM in HCMV-infected cells (Fig. 2D). HSF1-depleted, HCMV-infected monocytes also displayed increased sensitivity to cell death (Fig. 2F). Thus, these data demonstrate that elevated HSF1 activity following HCMV infection is necessary for the increased abundance of several pro-survival factors and the viability of infected monocyte.

**Figure 2:**
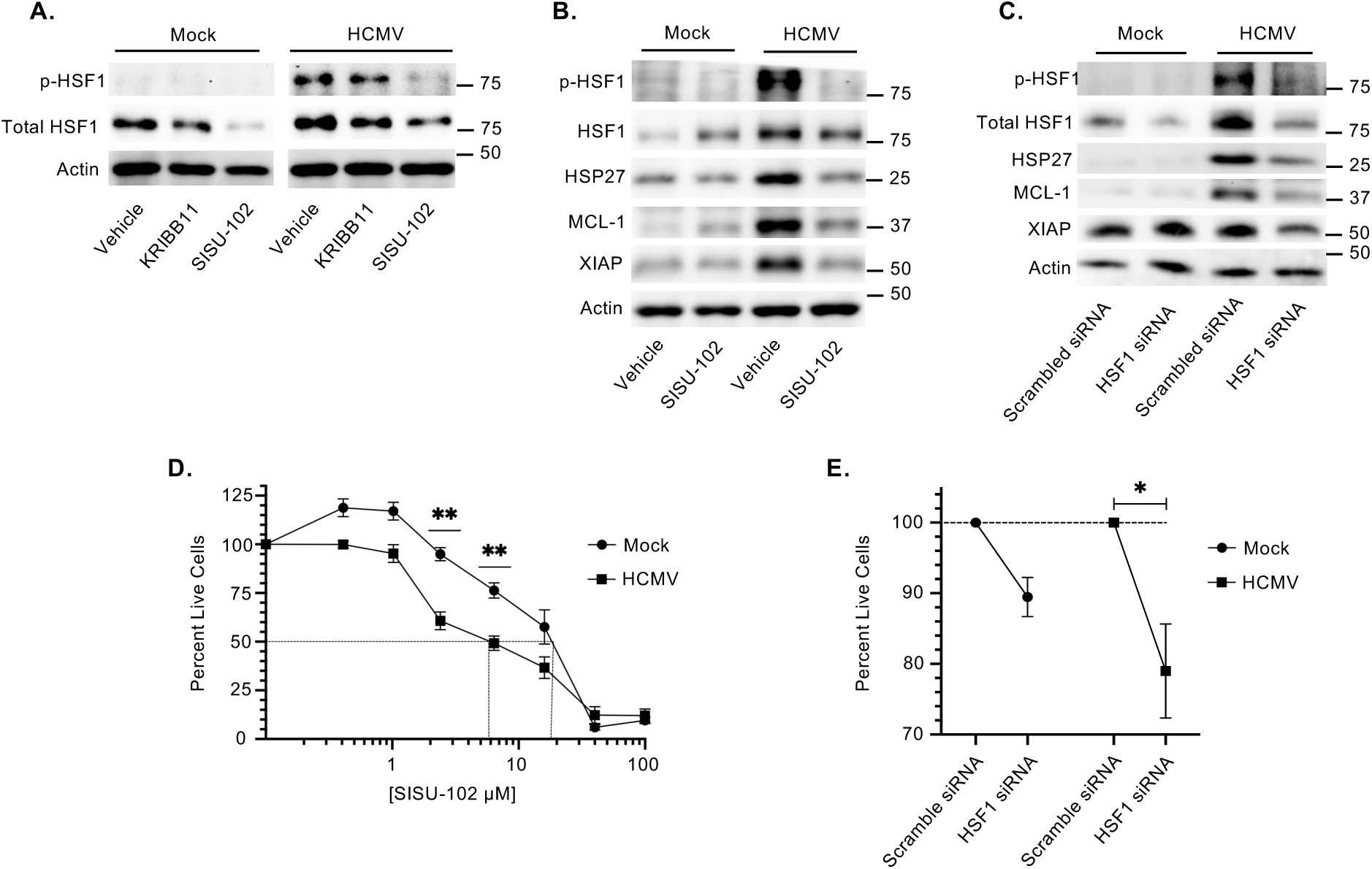
HSF1 is necessary for the survival of HCMV-infected monocytes. (A, B) Monocytes were pretreated with HSF1 inhibitors SISU-102 (5 μM) and K11 (5 μM) for 1 h. **(C)** Monocytes were transfected with a scrambled or HSF1-specific siRNA for 48 h. Following drug treatment or siRNA knockdown, cells were mock infected or infected (MOI 5) for an additional 24 h. Total HSF1, p-HSF1, HSP27, Mcl-1, and XIAP were detected by Western blot. Actin was used as a loading control. Western blots are representative of at least three biological replicates per group. **(D)** Monocytes were pretreated with increasing concentrations of SISU-102 for 1 h. **(E)** Monocytes were transfected with a scrambled or HSF1-specific siRNA for 48 h. Following drug treatment or siRNA knockdown, cells were mock- or HCMV-infected (MOI 5) for 24 h. Cell viability was assessed by annexin V and propidium iodide (PI) staining followed by flow cytometric analysis. All data are representative of at least 3 biological replicates per group. **P < 0.005 by one-way ANOVA with Tukey’s HSD post hoc test (D). *P < 0.05 by paired t-test (E).

### HCMV-induced mTOR activity requires HSF1

While HSF1 is primarily recognized as a stress-responsive transcription factor [67, 68], the absence of HSF1 binding sites within the promoters of several HCMV-induced pro-survival factors suggests alternative regulatory mechanisms for HSF1 aside from stimulating transcription. Beyond direct transcriptional activation, HSF1 has been implicated in the modulation of mTOR signaling [69, 70]. Consistent with our previous findings [34], we found that inhibition of HSF1 attenuates HCMV-induced mTOR activation with little effect on total mTOR levels, suggesting that HSF1 directly regulates the phosphorylation of mTOR (Fig. 3A). Accordingly, HSF1 is also necessary for the increased protein expression and phosphorylation of mTORC1 downstream substrates, including 4EBP-1 and S6K1, following HCMV infection. Parallel siRNA knockdown studies demonstrated similar requirement for HSF1 in mTORC1 signaling within infected monocytes (Fig. 3B). Upon a canonical stress response such as HS, HSF1 translocates to the nucleus to regulate gene expression [50, 71]. Yet, these data showing that HSF1 directly regulates mTOR phosphorylation suggest that HSF1 exerts a critical non-traditional function in protein translation in the cytoplasm of infected cells. In support, several studies have also demonstrated that HSF1 is not benign in the cytoplasm but can exert different biological activities dependent on its binding partners [49, 52, 72–75]. Thus, we sought to determine HSF1’s subcellular localization during HCMV infection. While HS treatment drove the expected translocation of HSF1 from the cytoplasm to the nucleus, HSF1 remained in the cytoplasm of HCMV-infected monocytes (Fig. 3C), despite being phosphorylated (Fig. 1A). Next, we examined whether cytoplasmic HSF1 interacts with mTOR. Co-immunoprecipitation (co-IP) assays in HCMV-infected monocytes demonstrate an interaction between HSF1 and mTOR, which was attenuated upon SISU-102 treatment (Fig. 3D and 3E). Because SISU-102 binds to the HSF1 DBD but only degrades nuclear and not cytoplasmic HSF1 [63], these data further suggest that SISU-102 can exert suppressive activity on cytoplasmic HSF1 during HCMV infection by blocking specific protein-protein interactions. Thus, HCMV-activated HSF1 is retained in the cytoplasm of HCMV-infected monocytes to engage and drive mTOR signaling.

**Figure 3:**
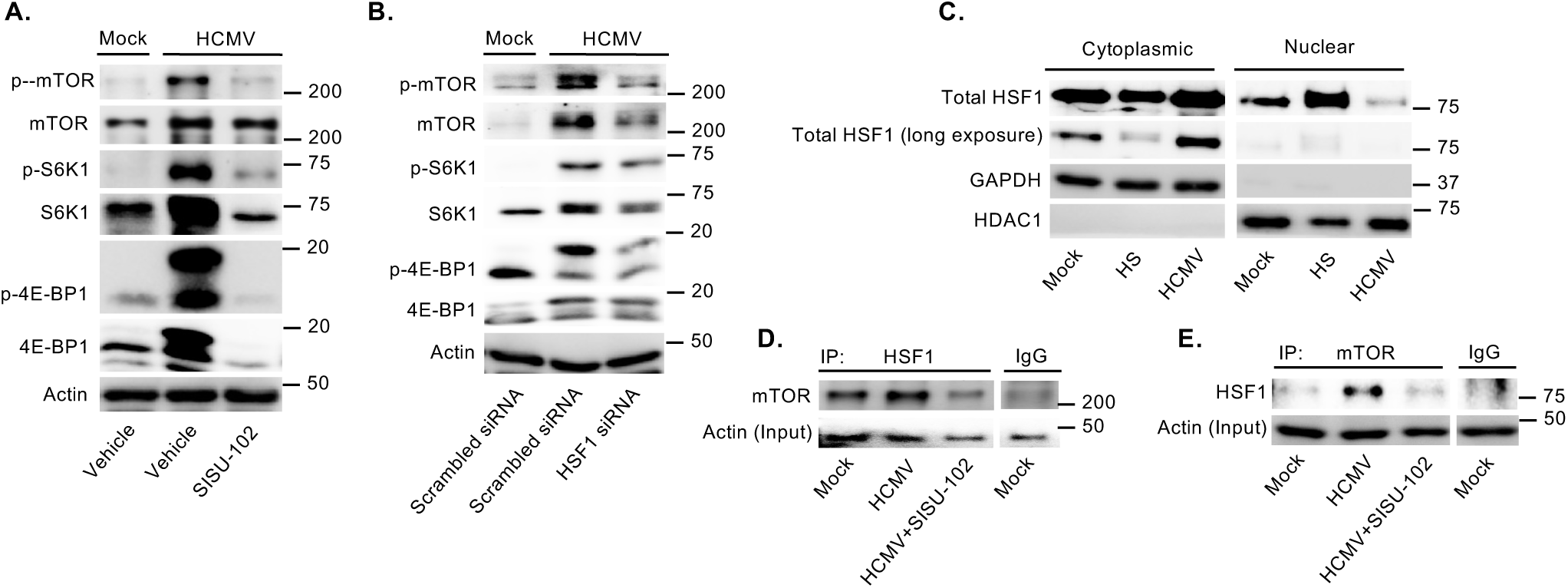
HCMV-induced mTOR activity requires HSF1. **(A)** Monocytes were pretreated with SISU-102 (5 μM) for 1 h. **(B)** Monocytes were transfected with a scrambled or HSF1-specific siRNA for 48 h. Following drug treatment or siRNA knockdown, cells were mock-infected or HCMV-infected (MOI 5) for an additional 24 h. Total mTOR, p-mTOR, S6K1, p-S6K1, 4E-BP1, and p4E-BP1 were detected by Western blot. Actin was used as a loading control. **(C)** Monocytes were mock- or HCMV-infected for 24 h (MOI 5). As a positive control, cells were subjected to HS (42°C) for 30 m and allowed to recover at 37°C for 24 h. Following treatments, nuclear/cytoplasmic fractionation was performed. Protein abundance of HSF1, GAPDH (cytoplasmic protein), and HDAC1 (nuclear protein) were detected by by Western blot. **(D, E)** Monocytes were mock- or HCMV-infected (MOI 5) for 16 h, followed by SISU102 (5 μM) treatment for 8 h. Co-immunoprecipitation was performed with anti-mTOR, anti-HSF1, or IgG isotype control antibodies. HSF1 and mTOR were detected by Western blot. Actin was used as an input loading control. All Western blots are representative of at least 3 biological replicates per group.

### HCMV-induced HSF1 stimulates global mRNA translation

During lytic replication, viral pUL38 functions to uncouple mTORC1-mediated mRNA translation from the canonical repressive effects of host stress responses [76]. However, pUL38 is not expressed in quiescently infected monocytes (Fig. 4A). As a control, infection with an HCMV mutant virus lacking US28 (ΔUS28) unable to establish quiescent infection led to pUL38 expression within monocytes. Together with our previous study demonstrating that HCMV induces mTORC1-mediated mRNA translation following infection of monocytes [36], these data suggest that HCMV employs a unique mechanism independent of viral lytic proteins to stimulate and maintain protein synthesis. Since activated HSF1 remains localized to the cytoplasm (Fig. 3C), where it interacts with mTOR and promotes downstream signaling within HCMV-infected monocytes, we asked whether HSF1 plays a non-canonical function in regulating mRNA translation. We utilized surface sensing of translation (SUnSET) assays, which exploit the incorporation of puromycin into elongating polypeptide chains, thereby terminating translation [77, 78], to investigate the impact of HSF1 on protein synthesis during infection. Consistent with previous studies demonstrating that HCMV stimulates global protein synthesis in infected cells [36], we found that HCMV increases mRNA translation by 24 hpi during the establishment of quiescence within monocytes (Fig. 4B & C). Puromycin incorporation into nascent peptides was reduced to mock levels by the presence of SISU-102, indicating the involvement of HSF1 during HCMV-driven translation. In contrast, the mechanism through which granulocyte-macrophage colony-stimulating factor (GM-CSF) stimulates cellular translation was independent of HSF1 (Fig. 4B & C). HSF1-depleted monocytes also exhibited decreased protein synthesis in response to HCMV infection, validating the involvement of HSF1 in regulating mRNA translation in infected monocytes (Fig. 4D & E). Further, HSF1-dependent translation following HCMV infection appears to be cell-type specific, as increased protein synthesis in lytically infected fibroblasts was unablated by the loss of HSF1 activity (Fig. 4F &G). Hence, activation of cytoplasmic HSF1 is necessary for the enhanced rates of mRNA translation within quiescently infected monocytes.

**Figure 4:**
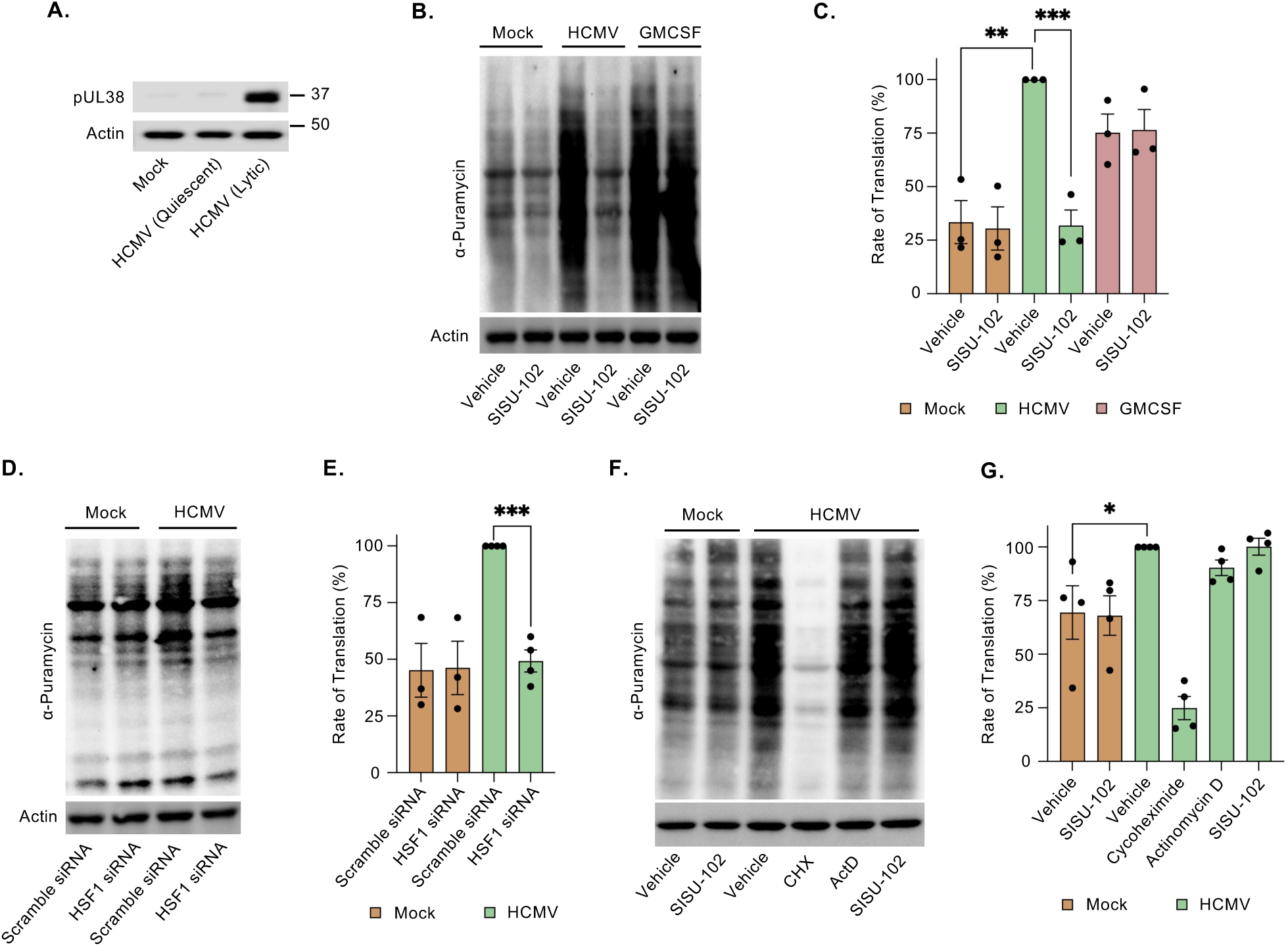
HCMV-induced HSF1 mediates a global increase of mRNA translation within infected monocytes. **(A)** Primary monocytes were infected with HCMV or ΔUS28 (MOI 5) for 24 h. pUL38 was detected by Western blot. Actin was used as a loading control. **(B, C)** Monocytes were mock- or HCMV-infected (MOI 5) or treated with GMCSF (100 ng/mL) for 24 h. DMSO or SISU102 (5 μM) was added at 23 h post infection or treatment for 1 h. **(D, E)** Monocytes were transfected with a scrambled or HSF1 siRNA for 24 h followed by mock or HCMV infection for an additional 24 h. **(F, G)** Confluent human embryonic lung 299 fibroblasts were mock or HCMV (MOI 5) infected for 24 h. DMSO, SISU102 (5 μM), cycloheximide (a protein synthesis inhibitor; 10 µg/mL), or actinomycin D (a transcription inhibitor; 10 μM) were added at 23 h for 1 h. Following treatments or siRNA knockdown, cells were pulsed with a low-dose of puromycin for 30 min to tag newly synthesized peptides. Puromycin labelling was detected by Western Blot. Actin was used as a loading control. **(C, E, G)** Densitometry was performed to measure total protein levels, which were normalized to actin. Western blots and densitometry are representative of at least three biological replicates per group. *P < 0.05, **P < 0.005, ***P < 0.005, by one-way ANOVA with Dunnett’s post hoc test.

### HSF1 and mTOR promote the translation of prosurvival transcripts

HCMV infection exerts a profound effect on the translatome of infected monocytes, distinct from the translational programs driven by canonical myeloid growth factors [36]. To define the HSF1- and mTOR-dependent translatome of HCMV-infected monocytes, we performed polyribosomal profiling on monocytes isolated from a single blood donor as an initial screen to globally identify translating mRNAs dependent on HSF1 and mTOR during HCMV infection. Although at least two biological replicates are generally performed with transcriptional and translational profiling studies, a single representative donor was selected because primary blood monocytes exhibit substantial donor variability regarding the magnitude of change in gene expression following HCMV infection. Thus, combining RNA-seq datasets from two donors can skew data to identify only genes that exhibit large expression changes across different donors, while subtle to moderate changes, which could have profound biological impact, are lost. However, to ensure the reproducibility of expression changes in genes of interest, we will validate these changes using a multitude of biochemical approaches with ≥3 independent blood donors. Analysis of the polyribosomal profiling dataset showed that HCMV infection upregulated 1,782 mRNAs by at least 2.5-fold within the polyribosome-associated pool compared to mock-infected monocytes (Fig. 5). Of these, 1,110 transcripts exhibited increased ribosome loading regardless of inhibitor treatment, suggesting that they are regulated independently of the HSF1-mTOR axis. We classified transcripts as HSF1- or mTOR-dependent if their ribosome association was reduced by >1.5-fold in the presence of SISU-102 or rapamycin, respectively. While only a small subset of transcripts showed a preferential dependence on ribosome loading on either mTOR alone (44 genes) or HSF1 alone (3 genes), the vast majority (625 genes) were dependent on both factors (Fig. 5 and supplemental Table 1). This high degree of overlap is consistent with our model, in which HSF1 directly interacts with and modulates mTOR activity to coordinate a specific translational program during infection.

**Figure 5:**
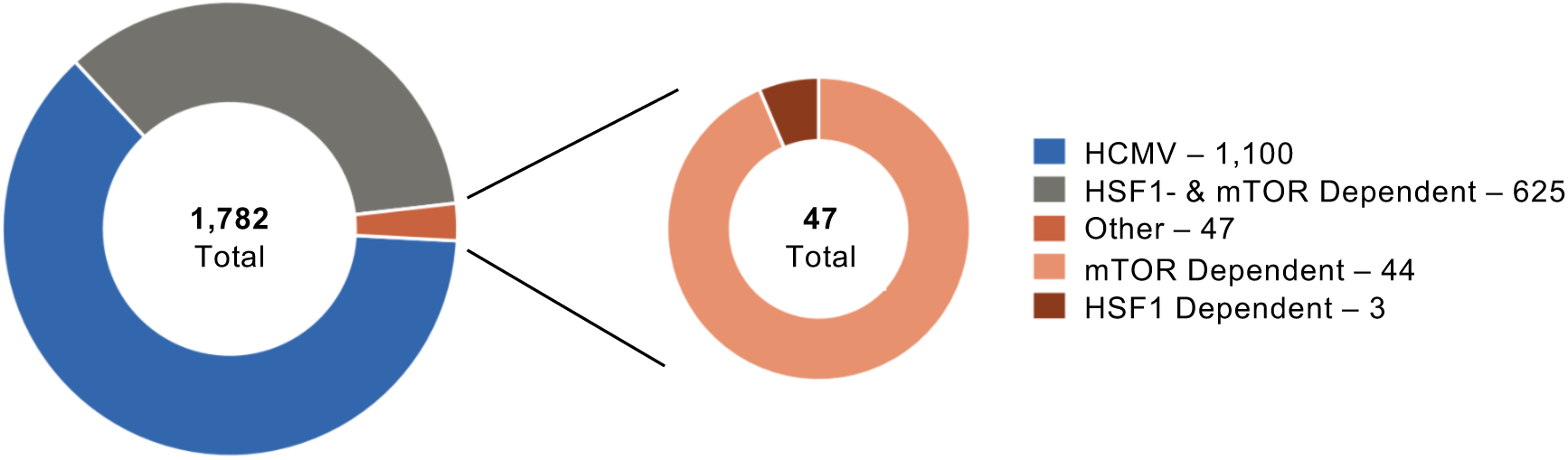
HSF1 and mTOR promote the translation of prosurvival transcripts following HCMV infection. Primary monocytes were pretreated with DMSO, rapamycin (5 μM), or SISU102 (5 μM) for 30 min. Cells were then mock- or HCMV-infected for 24 h (MOI 5) followed by polyribosomal profiling. HCMV-regulated genes were identified by being at least 2.5-fold over mock. HSF1 and/or mTOR dependent genes were reduced at least 1.5-fold in the presence of inhibitors.

### HCMV increases cIAP2 in a mTOR and HSF1-dependent manner to promote the survival of infected monocytes

In-depth analysis of HCMV-induced, ribosome-loaded transcripts identified cIAP2 (Cellular Inhibitor of Apoptosis Protein 2), which controls the processing of caspase-3 by limiting the conversion of the intermediate p19 subunit to the fully active the p17 subunit [79], as being dependent on both HSF1 and mTOR. As validation, we found that HCMV infection increases cIAP2 abundance and that inhibition of HSF1 or mTOR reduces protein levels within infected monocytes (Fig. 6A and 6B). To explore the role of the HSF1-mTOR signaling axis in regulating caspase-3 activation following HCMV infection of monocytes, we measured the abundance of procaspase-3 and its cleavage products (p19 and p17) following HCMV infection in the presence or absence of SISU-102 or rapamycin. As expected, HCMV infection resulted in the accumulation of procaspase-3 and p19 while limiting the generation of p17 (Fig. 6C). Indeed, pharmacological inhibition of either HSF1 or mTOR in infected cells led to the cleavage of procaspase-3 and the subsequent cleavage of p19 into the fully active p17. Next, HCMV-infected monocytes were treated with two structurally distinct cIAP2 inhibitors, birinapant or xevinapant, to assess the function of cIAP2 during HCMV-mediated suppression of caspase-3 activation. Similar to HSF1 and mTOR inhibition, blocking cIAP2 activity led to the cleavage of procaspase-3 and p19, resulting in the accumulation of p17 (Fig. 6D). Flow cytometric analysis of annexin V and PI staining showed a dose-dependent decrease in the viability of infected cells. Further, infected monocytes displayed increased sensitivity to cIAP2 inhibitors relative to uninfected cells, suggesting heightened role for cIAP2 in the survival of monocytes following HCMV infection (Fig. 6E). Collectively, these data establish the HSF1-mTOR signaling axis as a critical pathway in the rewiring of the host translational machinery required for the synthesis of cIAP2 and the survival of HCMV-infected monocytes.

**Figure 6:**
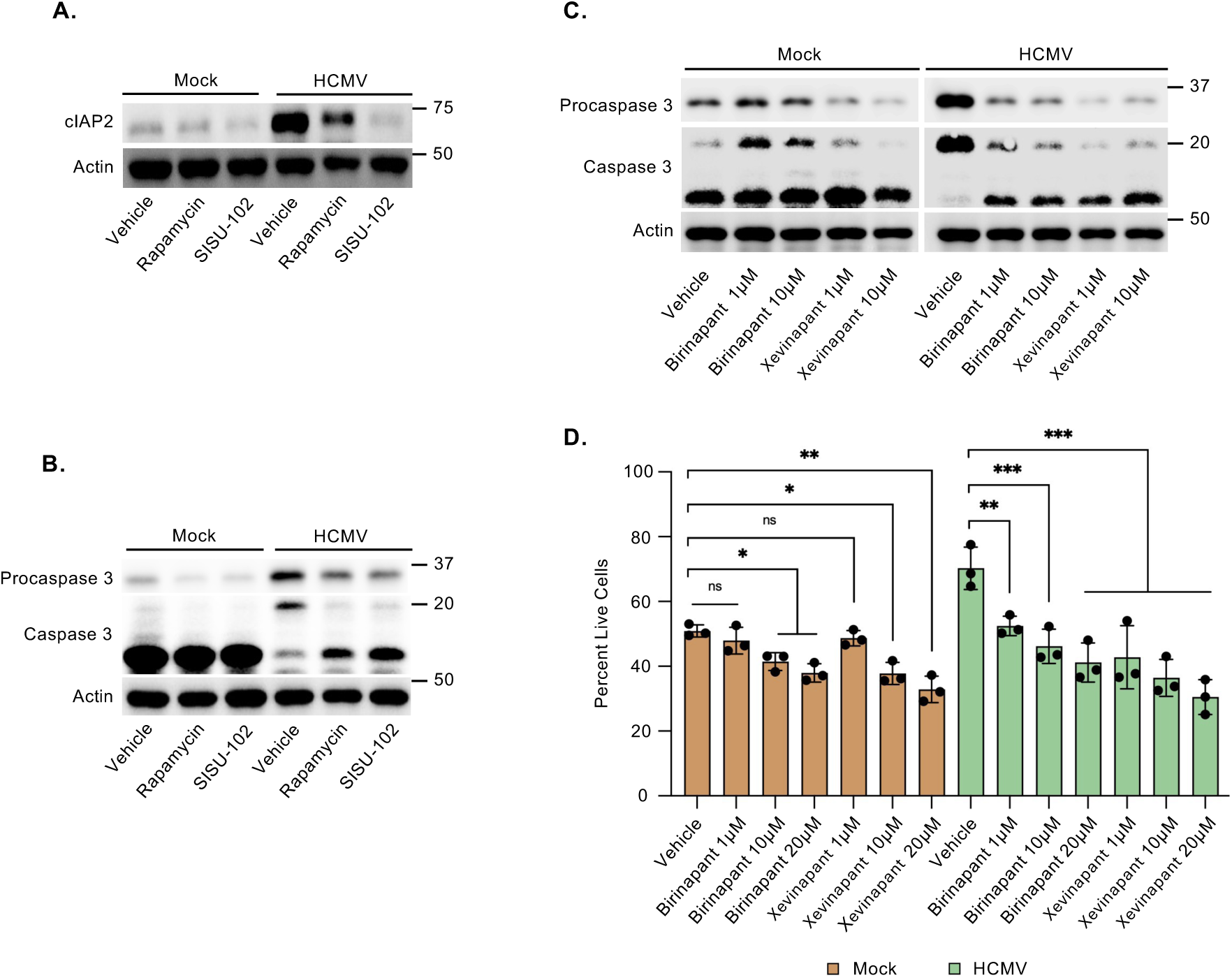
HCMV increases cIAP2 in a HSF1 and mTOR-dependent manner to promote the survival of infected monocytes. **(A, B)** Monocytes were pretreated with DMSO, rapamycin (5 μM), or SISU102 (5 μM) for 30 min. **(C, D)** Monocytes were pretreated with cIAP2 small-molecule inhibitors, birinapant or xevinapant, for 30 min. Following treatment with inhibitors, cells were then mock- or HCMV-infected for 24 h (MOI 5). **(A-C)** Abundance of cIAP2, procaspase-3, or caspase-3 were detected by Western blot. Actin was used as a loading control. Western blots are representative of at least three biological replicates per group. **(D)** Annexin V and PI staining was performed to assess viability by flow cytometry. Data are representative of ≥3 independent experiments. ns, not significant; *P < 0.05, **P < 0.005, ***P < 0.005, by one-way ANOVA with Dunnett’s post hoc test.

## DISCUSSION

Peripheral blood monocytes are considered the primary vehicles for the systemic dissemination of HCMV [20–24]. However, monocytes have a short 48-h lifespan and lack the permissivity for productive viral replication during this period [25]. To overcome these limitations, HCMV reshapes the host signaling environment to establish an antiapoptotic state, allowing for the differentiation of infected monocytes into replication-permissive macrophages [29–31]. Our previous work demonstrated that HCMV infection of monocytes globally upregulates protein translation through activation of mTORC1 and its downstream effectors [36]. In contrast, increased cellular translation by the normal myeloid growth factor GM-CSF is independent of mTORC1, indicating a distinct mechanism through which HCMV broadly modulates mRNA translation within infected monocytes [36]. In this study, we demonstrate that HCMV infection stimulates a non-canonical activation of cytoplasmic HSF1 to induce mTOR activity and drive a global cellular translational landscape directed towards a pro-survival state within infected monocytes.

The activation of HSF1 is a vital cellular response to maintain protein homeostasis and ensure overall cellular fitness during times of stress [49, 50]. However, our new data suggest that HSF1 activation following HCMV infection is not a classical host cell response to the virus, but a virally regulated event. HSF1 undergoes a rapid but transient phosphorylation under classical HS stress that resolves completely within 24 hours (Fig. 1A). In contrast, HCMV infection induces an early phosphorylation of HSF1 that is sustained through at least 24 hours. The contrasting activation kinetics of HSF1 between HCMV and HS suggest different mechanisms of activation between the two stimuli. Indeed, activation of HSF1 by HS is independent of the PI3K/Akt signaling pathway, while HCMV-induced HSF1 requires PI3K/Akt signaling (Fig. 1B). We have previously shown that the simultaneous engagement of HCMV glycoproteins gB and gH with EGFR and integrins, respectively [80], during viral entry stimulates a unique activation of Akt, leading to the increased abundance of a selected subset of Akt-dependent pro-survival proteins [32–34, 36]. It remains unclear how the virus-specific activation of Akt promotes the persistent activation of HSF1 following infection of monocytes without driving nuclear relocalization. Regardless, our data demonstrate that HCMV stimulates the continued activation of cytoplasmic HSF1, via the non-canonical activation of PI3K/Akt signaling, allowing for the maintenance of the antiapoptotic state and survival of infected monocytes.

Under homeostatic conditions, monomeric HSF1 is maintained in an inactive state in the cytoplasm through its association with different HSPs such as HSP70 and HSP90 [49, 51]. Upon HS or proteotoxic stress, accumulation of misfolded proteins recruits these chaperones away, releasing HSF1 and allowing it to trimerize and translocate into the nucleus [49]. We previously demonstrated that during lytic infection in fibroblasts, HCMV rapidly activates and induces the nuclear relocalization of HSF1 to promote viral gene expression [61]. In contrast, we observe that HCMV-activated HSF1 fails to localize to the nucleus during the infection of monocytes. This spatial regulation shows a fundamental divergence in how HCMV utilizes the HS response across different cell types. While nuclear translocation of HSF1 supports active, lytic replication in permissive cells [61], its retention in the cytoplasm in primary monocytes may be required to establish and maintain a quiescent infection. Future studies are key to addressing whether holding HSF1 within the cytoplasm of HCMV-infected monocytes is required for preventing lytic infection, and thus the establishment and maintenance of quiescent infection.

How HCMV activates HSF1 without inducing its relocalization into the nucleus during infection of monocytes remains unknown. We speculate that the nuclear translocation may require additional post-translational modifications such as secondary phosphorylation, acetylation, or sumoylation events triggered by HS but absent during HCMV infection. Without these secondary PTMs, the dissociation of HCMV-activated HSF1 from HSPs may be blocked resulting in the nuclear localization signal (NLS) being masked. Alternatively, chaperones locking HSF1 in the cytoplasm are also heavily regulated by PTMs [53]. Thus, HS and HCMV infection could also lead to distinct PTM patterns on HSP70 or HSP90 preventing chaperone dissociation. Regardless of the virus-specific mechanism regulating HSF1 nucleocytoplasmic shuttling, our data highlight the precision through which HCMV regulates the cellular HS response to drive functional changes conducive to long-term infection of monocytes.

Our data demonstrate that HCMV activates the cytoplasmic pool of HSF1 to engage mTOR, stimulating the translation of cellular pro-survival factors. However, the host energy-sensing kinase AMPK typically acts as a checkpoint to inhibit mTORC1 in response to stress stimuli shutting down cap-dependent translation to conserve resources [43–45]. During lytic replication in fibroblasts, HCMV effectively uncouples this metabolic checkpoint through the viral protein pUL38, which blocks AMPK and protects translation [46]. However, pUL38 is absent in quiescently infected monocytes (Fig. 4A). To bypass this limitation, we found cytoplasmic HSF1 functionally replaces pUL38 as the alternative factor that prevents AMPK-mediated repression of mTORC1 within infected monocytes by directly binding and activating mTOR (Fig, 3D, 3E). This targeted bypass allows for the translation of crucial pro-survival proteins like cIAP2, which limits caspase-3 activation and prevents host cell from apoptosis (Fig. 6A, 6C). Ultimately, by substituting the function of a lytic protein with a rewired host stress factor, HCMV transforms a short-lived monocyte into a long-lived cell responsible for facilitating systemic viral spread throughout the host.

These results carry significant implications for the development of future therapeutic interventions. Naturally, monocytes are short-lived cells and less than ideal for mediating the systemic dissemination of HCMV [25, 26, 28]. By activating HSF1 without stimulating nuclear translocation, HCMV can maintain cellular translation of cIAP2 and other pro-survival factors within infected monocytes, transforming them into long-lived “Trojan horses” able to deliver the virus to distant organ sites. Because HSF1 is also necessary for lytic replication [61], targeting HSF1 represents potential valuable therapeutic target capable of eliminating quiescently infected monocytes responsible for viral spread as well as preventing lytic replication at sites of infection. Further, as HSF1 is only activated in infected, stressed cells, targeting this stress factor could also minimize cytotoxic side effects to uninfected bystander cells.

## MATERIALS AND METHODS

### Human peripheral blood monocyte isolation

Human peripheral blood monocytes were isolated as previously described [30, 66, 81, 82]. Briefly, whole blood was obtained via venipuncture from random donors. All experimental protocols involving human subjects were conducted in accordance with the guidelines of the University Institutional Review Board (IRB) and the Health Insurance Portability and Accountability Act (HIPAA). Whole blood was diluted in RPMI 1640 medium (ATCC) and layered over Histopaque-1077 (MilliporeSigma) to eliminate erythrocytes and neutrophils. The remaining mononuclear fraction was collected, washed with saline to remove platelets, and further resolved by density gradient centrifugation through a Percoll gradient (40.48% and 47.70%; GE Healthcare). Purity of the isolated monocyte population was verified to be 90% via CD14- or CD16-positive surface marker staining. Isolated cells were washed with saline, resuspended in RPMI 1640, and counted. Unless otherwise specified, downstream experimental cultures were maintained in the absence of human serum at 37°C in a humidified 5% CO2 incubator.

### Virus preparation and infection

Human embryonic lung (HEL) 299 fibroblasts (CCL-137; ATCC) of low passage (P7–15) were cultured in Dulbecco’s Modified Eagle Medium (DMEM; Lonza) supplemented with 2.5 μg/mL plasmocin (Invivogen) and 10% fetal bovine serum (FBS; MilliporeSigma). For viral propagation, fibroblasts were infected with HCMV strain TB40/E. Upon observation of a 100% cytopathic effect, extracellular virus was harvested from the culture supernatant and cleared of cellular contaminants by ultracentrifugation (115,000 × *g*, 65 min, 22°C) through a 20% sorbitol cushion. The purified viral pellet was resuspended in RPMI 1640 medium (ATCC). For subsequent experiments, primary monocytes were infected at a multiplicity of infection (MOI) of 5 per cell (unless otherwise stated), which routinely yields an infection efficiency of >99% [18, 31]. Mock-infected controls were treated with an equivalent volume of virus-free RPMI 1640 medium.

### Western blot analysis

Monocytes were harvested in a modified radioimmunoprecipitation assay (RIPA) buffer (50 mM Tris-HCl [pH 7.5], 5 mM EDTA, 100 mM NaCl, 1% Triton X-100, 0.1% SDS, 10% glycerol) supplemented with protease inhibitor cocktail (MilliporeSigma) and phosphatase inhibitor cocktails 2 and 3 (MilliporeSigma) for 30 m on ice. Lysates were cleared of cell debris by centrifugation at 21,000 × *g* for 5 min at 4°C, and supernatants were stored at −20°C until further analysis. Protein samples were solubilized in Laemmli SDS sample nonreducing (6×) buffer (Boston Bioproducts) supplemented with β-mercaptoethanol (Amresco) by incubation at 95°C for 10 min. Equal amounts of total protein were loaded per lane, resolved via SDS-polyacrylamide gel electrophoresis (SDS-PAGE), and electroblotted onto polyvinylidene difluoride (PVDF) membranes (Bio-Rad). Membranes were blocked in 5% bovine serum albumin (BSA) (Fisher Scientific) for 1 h at room temperature and subsequently incubated overnight at 4°C with specific primary antibodies. The primary antibodies used were: α-puromycin (MilliporeSigma); α-phospho-HSF1 and α-HSF1 (Abcam); α-caspase 3 (Santa Cruz Biotechnology); α-p-mTOR (Santa Cruz Biotechnology); α-HSF1, α-phospho-mTOR, α-mTOR, α-phospho-4EBP-1, α-4EBP-1, α-Akt, α-phospho-Akt, α-S6K1, α-phospho-S6K1, α-cIAP2, α-GAPDH, α-Histone H3, and anti-HDAC1 (Cell Signaling Technology). Rhodamine α-β actin antibody (Bio-Rad) was used as loading control.

### SUnSET Assay peptide labelling

In vitro SUnSET assays were performed to measure global protein synthesis as previously described [77, 83]. Briefly, primary monocytes (3x10^6^ cells) or confluent HEL 299 fibroblasts (5 × 10^5^ cells) were cultured under the indicated experimental conditions. Following treatment, nascent polypeptide chains were pulse-labeled by adding puromycin (1 µM; MilliporeSigma) directly to the culture medium for 30 min prior to harvest. Cells were subsequently lysed, and the lysates were resolved via SDS-PAGE and transferred to membranes for Western blot analysis using an anti-puromycin monoclonal antibody. Total lane density was assessed with Bio-Rad’s Image Lab software to measure total protein levels.

### Cellular fractionation lysis gradient

Subcellular fractionation was performed using an iso-osmotic discontinuous iodixanol-based density gradient as previously described [84, 85], with minor modifications (87, 88). Briefly, primary monocytes (3 × 10^6^ monocytes/sample) were layered on top of a custom-prepared, iso-osmolar discontinuous iodixanol gradient (OptiPrep; MilliporeSigma). The gradient was centrifuged at 1,000 × *g* for 10 min at 4°C in a swinging bucket rotor. During centrifugation, monocytes migrated through a preliminary cell-wash layer before encountering a mild cell-lysis layer containing 0.5% IGEPAL CA-630 (MilliporeSigma). This detergent concentration selectively disrupts the plasma membrane while leaving the nuclear envelopes intact. Intact nuclei subsequently sedimented through a secondary wash layer and came to rest above a hyper-dense barrier (float layer). Soluble cytoplasmic fractions were harvested directly from the cell-lysis layer, whereas the crude nuclear fractions were isolated from the interface between the second wash layer and the underlying float layer. Both the cytoplasmic and nuclear fractions were subsequently processed and analyzed via Western blotting.

### Analysis of polysome-associated RNAs

Monocytes (10 × 10^7^ cells) were treated with 0.1 mg/ml cycloheximide (inhibitor of translation) at 37°C for 10 min prior to harvest. The cells were then washed with PBS containing cycloheximide at 4°C and pelleted by centrifugation at 4°C. Pellets were resuspended in polysome lysis buffer (20 mM Tris-HCl [pH 7.4], 140 mM KCl, 5 mM MgCl2, Triton X-100, 10 mM dithiothreitol [DTT], and 0.1 mg/ml CHX) and passed through a 27-gauge needle five times. Residual insoluble debris and mitochondria were pelleted by centrifugation for 10 m at 15,000 × *g* in a microcentrifuge. The clarified lysate was then layered onto a non-linear sucrose as described here [86]. Briefly, the non-linear sucrose gradient is comprised of three increasingly dense concentrations of sucrose (5%, 34%, and 55%). Polyribosome bound RNA cannot enter the hyper-dense 55% sucrose, resulting in the efficient collection of all polysome associated RNA into one fraction. The locations of ribosomal subunits, monosomes, and polysomes in the gradient were determined by continuous monitoring of the absorbance at an optical density of 254 nm (OD254) during gradient fractionation using a Brandel gradient fractionator system coupled to a spectrophotometer (ISCO). Total RNA was extracted from an equal volume of each gradient fraction using TRIzol, and contaminating DNA was removed with DNase (Thermo Fisher Scientific).

### RNA-Seq Analysis

RNAs were isolated out of TRIzol prior to library prep. RNA sequencing library preparation was preformed according to the illumina dual stranded mRNA prep kit. Library quality and QC metrics were assessed and conformed to adhere to Illumina’s standard using a bioanalyzer (Agilent Technologies). Completed RNA-Seq libraries were then sequences using the NextSeq system, and a 400 million read sequencing chip (Illumina). Sequences bases were then trimmed, aligned, and quantified to the HG38 human reference genome within the Partek flow Analysis suite. Following quantile normalization, differential gene expression analysis was conducted using Partek’s GSA (gene specific analysis) computational algorithm. Gene ontology and pathway analysis was conducted in the EnrichR analysis package. All graphs and figures were generated using Partek flow or various R-coding packages. The raw and processed data have been deposited in NCBI GEO (number will be provided upon final submission).

### siRNA silencing

Primary monocytes (3 × 10^6^ cells cells per transfection) were washed with phosphate-buffered saline (PBS) and resuspended in 100 μL of P3 Primary Cell Nucleofector Solution (Lonza). Cells were mixed with 100 nM Silencer Select HSF1 targeting siRNA (Ambion-Thermo Fisher Scientific) or 100 nM Silencer Select scramble control siRNA (Ambion-Thermo Fisher Scientific). Transfection was executed using a 4D-Nucleofector System (Lonza) under program EI-100. Following nucleofection, monocytes were immediately transferred to RPMI 1640 medium supplemented with 2% human AB serum and allowed to recover for 24 h at 37°C. The transfected cells were subsequently mock-infected or infected with HCMV for an additional 24 h and subjected to immunoblot or flow cytometry analysis.

### Flow Cytometry

Primary monocytes were washed in PBS and incubated in blocking solution consisting of fluorescence-activated cell sorting (FACS) buffer (145 mM NaCl, 8.45 mM Na2HPO4, 1.83 mMNaH2PO4, and 0.1% NaN3), 5% BSA, and human FcR-binding inhibitor (eBioscience) for 20 min on ice. Following Fc-receptor blocking, cells were stained on ice with an allophycocyanin (APC)-conjugated anti-CD14 antibody or an APC-conjugated mouse IgG1 isotype control (both from BioLegend). To identify apoptotic and necrotic populations, cells were washed and subsequently stained with fluorescein isothiocyanate (FITC)-Annexin V and propidium iodide (PI) (Thermo Fisher Scientific). Cells were then analyzed by flowcytometry using an LSRFortessa cell analyzer and BD FlowJo software (BD Biosciences). Double-negative cells represent live cells, whereas double- and single-positive cells represent dead and/or dying cells.

### Statistical Analysis

All experiments were performed with a minimum of three independent biological replicates using primary cells isolated from distinct blood donors. Data were analyzed with GraphPad Prism (San Diego, CA) software using Student’s t test or one-way ANOVA with multiple comparisons when appropriate. *P*-values less than 0.05 were considered statistically significant.

## Acknowledgments

We thank Wayne Decatur and Christine Burrer in the Department of Microbiology and Immunology at SUNY Upstate Medical University for technical support, maintenance of lab operations, and assistance with virus growth and isolation. National Institute of Allergy and Infectious Disease (R01 AI170834 and R01 AI141460) to G.C. Chan, and National Heart, Lung, and Blood Institute (R01 HL139824) to G.C. Chan.

